# Effects of optogenetic silencing the Anterior Cingulate Cortex in a delayed non-match to trajectory task

**DOI:** 10.1101/2023.08.30.555490

**Authors:** Ana S Cruz, Sara Cruz, Miguel Remondes

## Abstract

Working memory is a fundamental cognitive ability, allowing us to keep information in memory for the time needed to perform a given task. A complex neural circuit fulfills these functions, among which is the anterior cingulate cortex (CG). Functionally and anatomically connected to the medial prefrontal, retrosplenial, midcingulate and hippocampus, as well as motor cortices, CG has been implicated in retrieving appropriate information when needed to select and control appropriate behavior. The role of cingulate cortex in working memory-guided behaviors remains unclear due to the lack of studies reversibly interfering with its activity during specific epochs of working memory. We used light to activate eNpHR3.0 and thus silence cingulate neurons while animals perform a standard delayed non-match to trajectory task. We found that, while not causing an impairment in eNpHR3.0+ animals when compared to eNpHR3.0-CTRL animals, silencing cingulate neurons during retrieval decreases the mean performance if compared to silencing during encoding. Such retrieval-associated changes are accompanied by longer delays in control animals, consistent with an adaptive recruitment of additional cognitive resources.

## Introduction

Working memory (WM), our ability to hold retrievable task-relevant information in memory for the time necessary to perform the task, plays a significant role in our daily life (Draschkow et al., 2021). In both primates and rodents, spatial WM relies on the top-down executive control of the overlapping anterior cingulate cortex (CG) and medial prefrontal cortex (mPFC) divisions (Battaglia et al., 2011; Jones and Wilson, 2005; Li et al., 2021; Rajasethupathy et al., 2015) responsible for selecting and retrieving the appropriate information when needed.

CG neural activity has been associated with fundamental cognitive functions in primates, namely executive functions supporting cognitive control (Botvinick et al., 2004), encoding and associating motivational context by mapping positive and negative outcomes to actions during decision-making (Hayden and Platt, 2010; Kennerley et al., 2006), processing aversive stimuli (Johansen et al., 2001), detecting surprise (Haddon and Killcross, 2006; Inokawa et al., 2010), conflict (Pardo et al., 1990) and errors (Carter et al., 1998). In rodents, the CG region has been associated with mnemonic functions involving the consolidation of motivationally salient contextual cues (Frankland et al., 2004; Goshen et al., 2011; Hoi et al., 2008; Kol et al., 2020; Weible et al., 2012), and sequential decision-making based on the retrieval of memorized associations between context, effort and reward (Akam et al., 2021; Fatahi et al., 2018; Porter et al., 2019; Remondes and Wilson, 2013). Generally, across species, CG is considered crucial for associating actions to outcomes in their episodic context, signaling errors, and suppressing inappropriate responses, all fundamental components of working memory (Nyberg and Eriksson, 2016).

During spatial working memory tasks the activity of CG neurons reflects available spatial trajectories (Remondes and Wilson, 2013), and exhibits complex, finely tuned interactions with the hippocampus (HIPP), where spatial information is stored (Burgess et al., 2002; O’Keefe, 1993). Moreover, CG neurons’ activity accompanies the retrieval of contextual memory items from CA3-CA1 HIPP neural circuits (Rajasethupathy et al., 2015; Remondes and Wilson, 2015). The role CG neural activity might play in spatial working memory has never been tested beyond permanent lesions, and poorly controlled silencing using non-specific local drug infusions, yielding conflicting results (Gisquet-Verrier and Delatour, 2000). In a spatial working memory version of the Morris Water Maze, near-complete aspirations of both CG and adjacent anterior midcingulate cortices (MCC), resulted in learning and memory impairments compensated by extra training, contrary to lesions of both posterior MCC and retrosplenial (RSC) cortices, whose resulting deficits where permanent (Sutherland et al., 1988). CG-specific lesions result in very clear working memory deficits in a go-no-go task, accompanied by the absence thereof after mPFC lesions (Gisquet-Verrier and Delatour, 2000), mild to absent effects in a delayed-alternation task unless complete mPFC ablation was performed (Delatour and Gisquet-Verrier, 2001; Fritts et al., 1998; Shaw and Aggleton, 1993), or increased errors exclusive to the learning phase of the task (Sánchez-Santed et al., 1997). This is also reported for another WM task, the 8-arm radial maze (Joel et al., 1997), with only combined mPFC lesions resulting in deficits, something replicated in a WM version of the Morris Water Maze (Sloan et al., 2006). These and other WM tasks require animals to retrieve the memory of a previous decision, and of the time it occurred, to disambiguate it from similar, previously rewarded, decisions (Aggleton et al., 1986; Maguire, 2001; Shaw and Aggleton, 1995; Shaw et al., 2013; Wilson et al., 2013). Likewise, the most widely used WM task, the Delayed Non-Match to Place task (DNMP), requires an animal to experience a delay period between a “Sample”, or encoding, run, in which one of the choice arms is blocked and animals are rewarded for coursing the open arm, and a “Test”, or retrieval, run in which both arms are made available, with animals rewarded for choosing the arm previously blocked. In a “Match-to-Trajectory” (DMT) version of this task, in which rats must match a previous trajectory on a T-shaped maze, CG-specific lesions resulted in deficits, with increased perseverance (Dias and Aggleton, 2000), contrary to their spontaneous alternating behavior (Richman et al., 1986), and increased errors after 10 and 20 sec delays between Sample and Test (Ragozzino and Kesner, 2001).

Given the above data, whether the anterior cingulate cortex is necessary for spatial working memory remains an open question, and several hypothetical scenarios are possible. While CG might be uniquely necessary for specific information processing that does not occur outside its limits, it is also quite plausible that the long-lasting to permanent nature of infusions or lesions triggers an adaptive process, in which supplementary motor areas, neighboring cortices MCC, RSC, or even HIPP take over WM-related functions, given the extensive mutual anatomical and functional connectivity (Ferreira-Fernandes et al., 2019), or simply compensate with other cognitive mechanisms. The fact that in the above studies CG is permanently affected, across all trials within one session, precludes any direct causal attribution of performance deficits to CG neural activity, when WM is manifested in individual trials. Such causal attribution warrants experiments that interleave, in the same session, trials in which CG neurons are silenced, or not, at distinct epochs.

Such types of manipulation became possible with the development of optogenetics. Here, light-activated membrane-bound molecules with the capacity to sustainedly, and reversibly, change neuronal membrane potential, either by opening ion channels or pumping ions across the membrane (L Fenno, 2010), allowed the possibility of silencing groups of neurons with behavior-specific temporal accuracy. Indeed, optogenetics is the only technique allowing for reversible perturbation of neural activity within the time constants of the cognitive functions at play during working memory (Yamamoto et al., 2014). Of the optogenetic neuronal silencers available, eNpHR3.0, an enhanced version of an earlier halo-rhodopsin, effectively hyperpolarizes neurons in vitro for over 5 seconds (Takeuchi et al., 2022; Zhang et al., 2019), enough to suppress action potentials in expressing neurons for up to 20 sec, with diverse behavioral effects in distinct brain regions, (Ferenczi et al., 2016; Imayoshi et al., 2020; Kang and Han, 2021; Mazzitelli et al., 2021; Moon et al., 2017; Van den Oever et al., 2013; Orsini et al., 2017; Pluta et al., 2019; Tye et al., 2011; Zhuang et al., 2021), which makes it ideal for the silencing of CG neurons during DNMT.

To test the hypothesis that CG is necessary for WM, once the task is learned, and to try and detangle its putative role in WM encoding versus retrieval, we used viral vectors to express the optogenetic silencer eNpHR3.0 in all CG neurons, by locally injecting either AAV9-hSyn-eNpHR3.0-eYFP or an “empty” CTRL construct AAV9-hSyn-eYFP, in 10 and 7 adult male Long Evans rats, respectively, subsequently trained in a DNMT task to 75% performance, and ran multiple 30-trial DNMT sessions, in which we interleaved three types of trials: no-illumination (NI), illumination during the Test, or retrieval, phase (TI), and illumination during the Sample, or encoding, phase (SI), with Orange (620 nm) light delivered via fiberoptic implants. A delay of 30 seconds was imposed between Sample and Test runs.

For each animal we measured three performance variables: individual trial and global performance (% correct trials), response latency to the choice point (CP), and time spent at the CP. We then used multivariable Generalized Linear Mixed Model (GLMM) using the above categorical and continuous variables, to investigate the impact of silencing CG neurons in WM-dependent behavior.

## Results

### Running the DNMT protocol with fiberoptic stubs and patch cords attached does not change behavioral performance

Freely moving behavior is conditioned by the relations of body shape and movement, with surrounding context structure and dynamics. As such, it is imperative that pre-manipulation WM testing is run under the same conditions as those used during CG silencing, except for light delivery. We thus ran the DNMT protocol in two groups of animals expressing either AAV9-hSyn-eNpHR3.0-eYFP (eNpHR3.0+, 10 rats) or an empty construct (CTRL, 7 rats) AAV9-hSyn-eYFP (Figure 1A-B and S1). We found no impairments in DNMT performance, nor significant differences between the two groups during this pre-illumination phase (Figure 2 A-B, t-test, t=1.45, p= 0.153) suggesting that, prior to the optogenetic activation protocol, all groups exhibited similar levels of task proficiency. We also found no significant differences between the groups’ mean latencies to reach the choice point during the pre-illumination phase (Figure 2C, Mann-Whitney, U=172802.5, p=0.75. CTRL: N= 545 trials, 2.811 s. eNpHR3.0+: N= 641 trials, 3.010 s, median). In both groups, rats tend to take longer to reach the choice point in error trials, with no significant difference between eNpHR3.0+ and CTRL groups (Figure 2D, Mann-Whitney, U=1, p=1.0. CTRL: N= 99 error trials. 3.170 s. eNpHR3.0+: N= 100 error trials, 2.744 s, median). Likewise, we computed the time spent at the CP and found no significant differences between the groups during the pre-illumination period (Figure 2E, Mann-Whitney, U=169897.5, p=0.417. CTRL: N= 545 trials, 0.893 s. eNpHR3.0+: N= 641 trials, 0.893 s. median). Like before, rats spend longer at the CP in error trials, with no significant difference between groups (Figure 2F, Mann-Whitney, U=0.0, p=1.0. CTRL: N= 99, 0.992 s. eNpHR3.0+: N= 100, 1.12 s, median).

**Figure 1.**
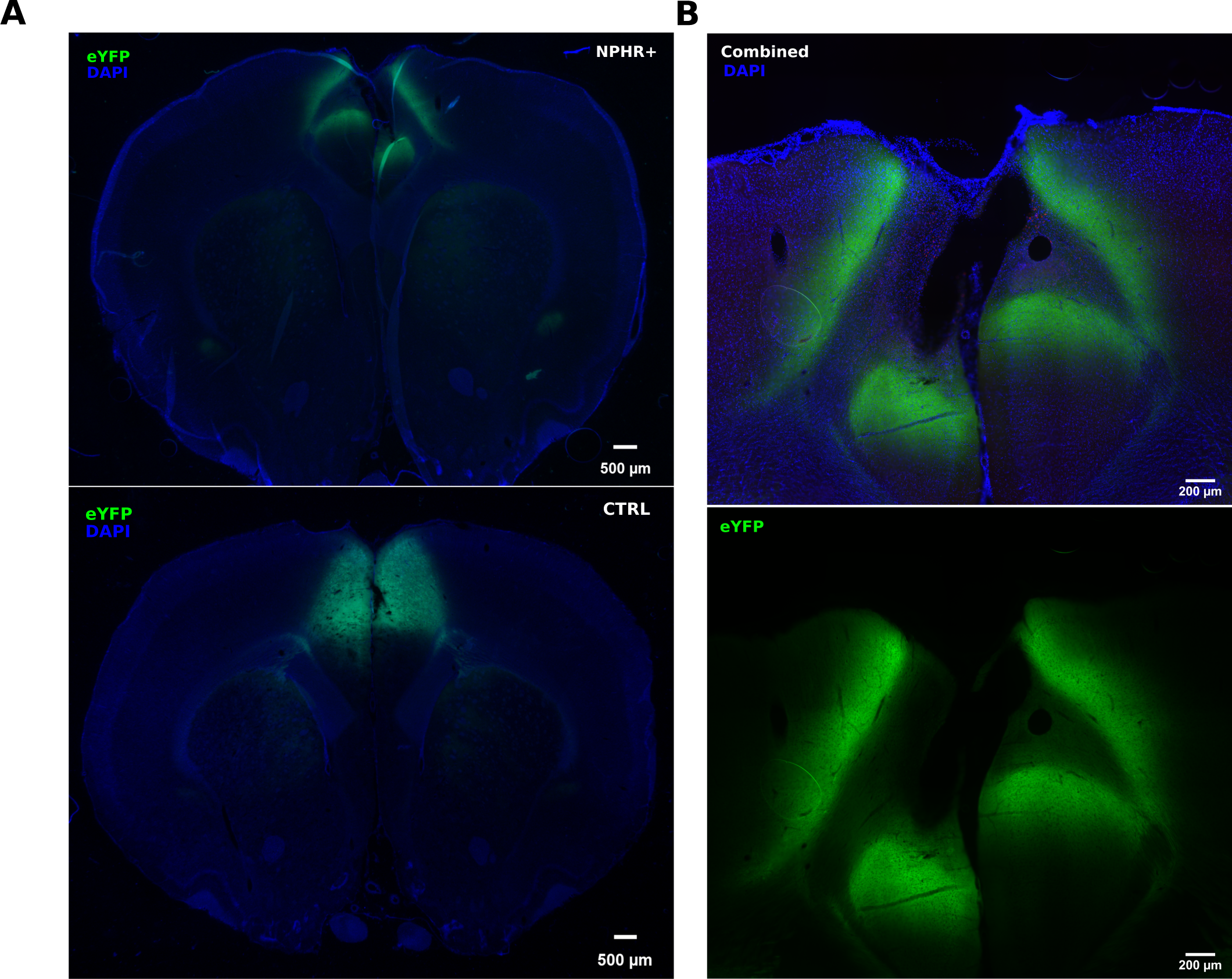
Anatomical distribution of eYFP fluorescence expression in the anterior cingulate cortex. (A) Expression of eYFP (top) in an eNpHR3.0+ and eNpHR3.0-(CTRL) rat (bottom) (B) Higher magnification of the three imaged channels on another example of an eNpHR3.0+ slice+.

**Figure 2.**
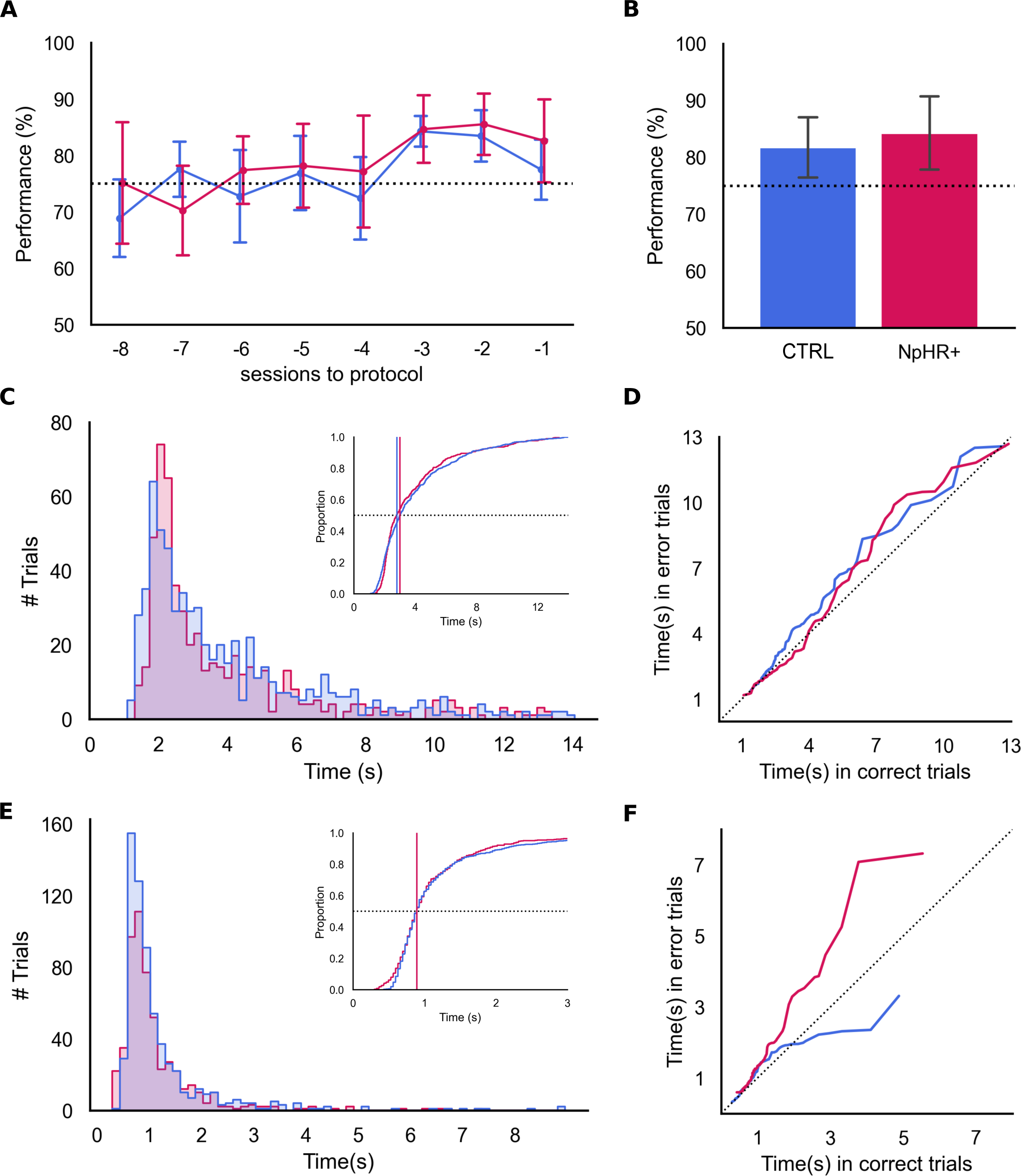
Performance during baseline sessions. Before illumination sessions begin there are no differences between experimental groups. (A) Group performances in the ten sessions preceding the illumination protocol onset and (B) in baseline sessions. We found no significant differences between group performances (t-test, t=1.451, p=0.153, CTRL: N= 7, 81.73% ± 5.44, eNpHR3.0+: N=10, 84.26% ± 6.55, mean ± sd). (C) Distributions of the latency to reach the choice point, in seconds, for both groups during baseline sessions. No significant difference between group medians was found (Mann-Whitney, U=17282.5, p=0.750, CTRL: N= 545 trials, 2.811 s. eNpHR3.0+: N= 641 trials, 3.009 s, median). (D) Quantile-quantile baseline latency to choice point distribution comparison between correct and error trials for both groups. We found no significant difference between group medians for baseline error trials (Mann-Whitney, U=1, p=1. CTRL: N= 99 trials, 3.170 s. eNpHR3.0+: N= 100 trials, 2.744 s, median). (E) Similar to C but for the time spent at the choice point, in seconds. No significant difference between medians was found (Mann-Whitney, U=169897.5, p=0.417. CTRL: N= 99, 0.893 s. eNpHR3.0+: N= 100, 0.893 s). (F) Similar to (D), for the time spent at the choice point. Once more, we found no significant difference between the medians of baseline error trials when comparing both groups (Mann-Whitney, U=0, p=1. CTRL: N=99, 0.992 s. eNpHR3.0+: N=100, 1.124).

### Light delivery during DNMT results in similar performances between CTRL and eNpHR3.0 animals

After the “baseline” phase above, we started the “illumination” phase protocol, in which “Test-illuminated” trials (TI), with light delivered through the CG fiberoptic implants in the Test epoch of the DNMT protocol, were randomly interleaved with “Sample-illuminated” trials (SI), in which light delivery was restricted to the Sample epoch, and with “non-illuminated” trials (NI), where no light was delivered, to account for individual animal variability (Fig. 3 A-D).

**Figure 3.**
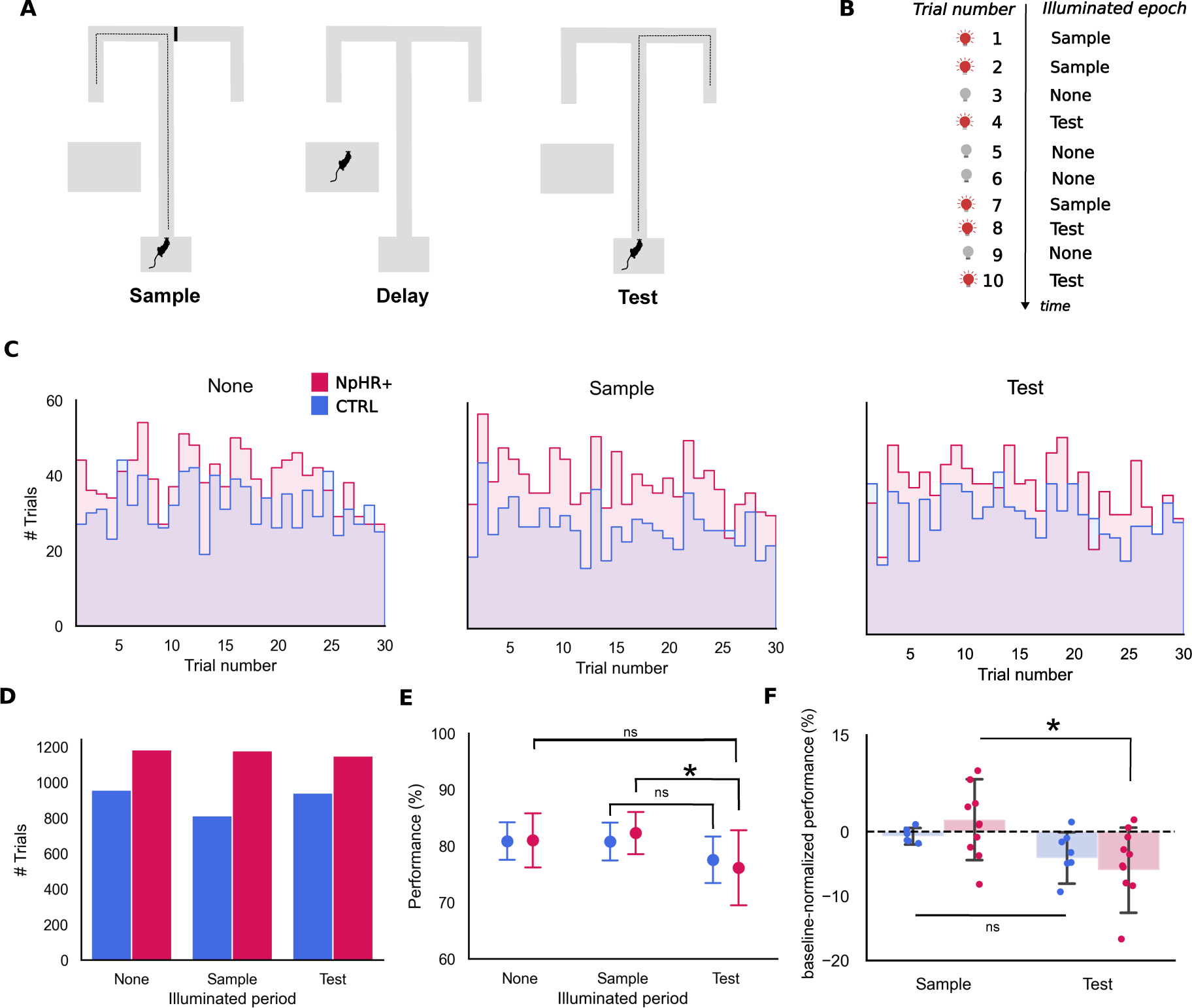
Performance and trial distributions during Illumination sessions. (A). Delayed non-match to position task. In this task the rat runs a randomly assigned forced trajectory in the sample epoch, followed by a delay of 30 seconds, after which it must follow a non-match rule to collect a reward at the end of the chosen trajectory. The ITI (inter-trial interval) ranged between 20 to 30 seconds. (B) Schematic representation of a sequence of trials according to illumination epoch (None, Sample, Test). Illumination epoch was randomly assigned a trial number within the session, with no more than 3 consecutive trials of the same type. (C) Distribution of manipulation types (None, Sample, Test) across the fixed structure of sessions (30 trials per session) for each experimental group (CTRL in blue, eNpHR3.0+ in red). (D) Total numbers of the three manipulation types. No-illumination (NI, “None”: CTRL: 958 trials. eNpHR3.0+: 1184 trials), sample illumination (SI, “Sample”, CTRL: 813 trials. eNpHR3.0+: 1178 trials) and test illumination (TI, “Test”, CTRL: 940 trials. eNpHR3.0+: 1150 trials). Trials in which the rat did not express a choice within 15 seconds after departure were excluded from the analysis. (E) Group performances (% correct) for each manipulation type and animal group, mean+/−SD, (CTRL: None: 80.88±3.62, Sample: 80.80±3.70, Test: 77.54±4.45. eNpHR3.0+: None: 81.00±5.08, Sample: 82.30±3.97, Test: 76.15±7.01). (F) NI-normalized performance in Test vs Sample in both animal groups. The bars represent the average difference (mean+/−SD), and the dots represent individual performance differences (illuminated - non-illuminated) (CTRL: Sample: −0.58±1.14 %. Test: - 3.34±3.48 %. eNpHR3.0+: Sample: 1.29±5.40 % Test: −4.85±5.38 %). All summary data depicted is mean+/−SD.

Overall, we observed no performance differences when directly comparing the two groups of animals, CTRL vs eNpHR3.0, indicating that eNpHR3.0 activation in CG does not absolutely impair decision making in the DNMT task. However, when we compare trials in which no light was delivered (NI) with either Sample-illuminated (SI) or Test-illuminated (TI) ones, we found a significant effect of the illumination phase on trial outcome (Fig. 3 E, X^2^=11.66192, p=0.003, Fixed Effect Omnibus test). Importantly, *post-hoc* analyses revealed that this TI-related decreased performance was restricted to the eNpHR3.0+ group (p=0.032), with more errors committed when CG was illuminated in Test than in Sample trials. The observed performance difference does not extend to CTRL animals.

We observed considerable variation in performance between animals, within each group. We sought to minimize its effects by normalizing (through subtraction) each animaĺs performance in TI and SI by their performance in NI, when no light is delivered. This analysis further clarified the difference we found when comparing sample vs test-illuminated performance, as well as its specificity to eNpHR3.0+ animals (Figure 3 E-F). The fact that light-activation of eNpHR3.0 is associated with decreased performance in TI compared SI, suggests that, while not causing an impairment if compared to empty-vector CTRL animals, within the eNpHR3.0+ group, CG neuron silencing during retrieval leads to lower performance levels than the same manipulation during encoding.

### In CTRL, but not eNpHR3.0+ animals, latency to reach the choice point is increased when light is delivered during retrieval

To look deeper into the dynamic of the decision process we measured the temporal delays in pre-defined trial periods, latency between trial start and choice point (CP), and time spent at the CP. While we found no differences between the two groups of animals, CTRL vs eNpHR3.0, in the overall latency, we found a significant effect of the illumination condition on latency to reach the CP (Figure 4A-C, X^2^=12.4618, p=0.002, Fixed Effect Omnibus test), with CTRL animals exhibiting an increased delay whenever light is delivered during Test (Fig 4B, 100-200ms, p=0.04 *Post Hoc* comparisons). The lengthier trials seen within the CTRL group in TI vs SI might be the result of a distraction, or a light-mediated direct effect on neurons (Owen et al., 2019; Stujenske et al., 2015), one that CTRL animals manage to overcome, by virtue of recruiting additional, time-consuming, cognitive resources, made unavailable in eNpHR3.0+ animals. Thus, the fact that no such difference is found in eNpHR3.0+ animals, even though retrieval is also presumably interrupted, reflects the lack of such additional resources, which results in no extra time consumed for its implementation.

**Figure 4.**
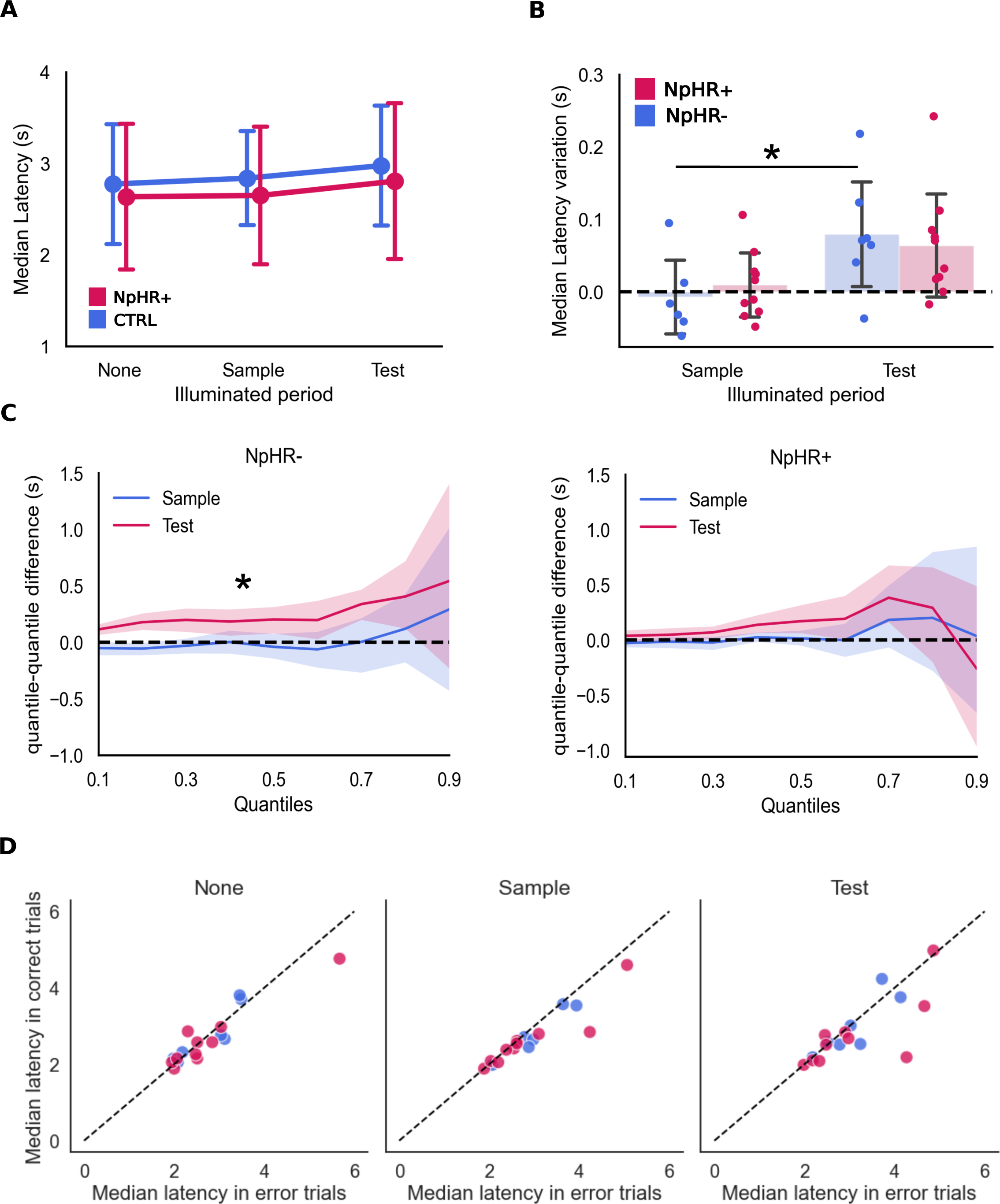
Latency to reach the choice point. (A). Latency to choice point medians across groups and manipulation types (CTRL: None: 2.77± 0.71 s. Sample:2.84±0.56 s. Test: 2.98±0.71 vs. eNpHR3.0+: None: 2.64±0.84 s. Sample: 2.65±0.79 s. Test: 2.81±0.90 s, mean ± sd). (B) Latency variation from non-illuminated trials, in seconds. The bars represent the average within-animal median difference (mean+/−SD), and the dots represent median latency differences within each animal (median illuminated - median non-illuminated) (CTRL: Sample: −0.007±0.056 s. Test:0.079±0.078 s. eNpHR3.0+: Sample: 0.009±0.047 s Test: 0.064±0.074 s). (C) Similar to (B) but across all quantiles of the distributions, in seconds, for the sample illuminated trials (left) and test illuminated trials (right) (mean± CI(95%)) (D) Comparison between the median latencies to reach the choice point in error vs. correct trials. Each dot corresponds to one rat. The dashed line corresponds to a situation in which both medians (in correct and error trials) have the same value. All summary data depicted is mean+/−SD.

## Discussion

A major obstacle in dissecting the neural bases of memory-guided behavior lies in reconciling multiple studies describing the effects of long-lasting silencing and lesions. Here we used optogenetics to silence CG neurons bilaterally in rats pre-trained to 75% criterium in a DNMT task, at distinct phases of individual trials, and found that activating eNpHR3.0 to silence CG does not result in significant changes in performance relative to the effects of light delivery in CTRL animals. However, in eNpHR3.0+, but not CTRL, animals, we found a significant effect of light delivery whereupon decreased performance is observed whenever CG is illuminated with 620 nm light during “Test”, presumably memory retrieval, if compared to the one observed following illumination during Sample (encoding). Conversely, the median time eNpHR3.0+ animals take to reach the choice point is unaffected by illumination, but is increased in CTRL animals when light delivery occurs during Test, indicating a slower decision process. The nature of the observed effects suggests that, in eNpHR3.0+ animals, CG opsin activation in neurons adds its effects to an already existing light-mediated interference with neural activity. Light alone can decrease firing rate in striatal medium spiny neurons and produce behavioral effects due to the activation of an inwardly rectifying potassium conductance (Owen et al., 2019), especially under prolonged illumination conditions. Furthermore, 5-10 mW illumination of CG neurons with 532 nm light resulted in a 30-40% increase in firing rate (Stujenske et al., 2015). Besides a specific role of CG neural activity in working memory, the observed effects might also be explained by an indirect effect of our manipulation, involving visual attentional control (Koike and al., 2015), anticipatory, corrective, attention, causing an increase in performance after sample illumination and resulting in significant differences between SI and TI in eNpHR3.0+ animals (Norman et al., 2021), and/or affecting sustained attention at any point of the task (Jendryka et al., 2023).

The lengthier trials seen within the CTRL group in TI vs SI (Fig 4C) might represent the recruitment of additional cognitive resources to face light-mediated changes, (Owen et al., 2019; Stujenske et al., 2015), allowing them to overcome this effect, unlike eNpHR3.0+ animals.

In sum, our data shows that using eNpHR3.0 for optogenetic silencing CG neurons does not result in absolute working memory impairments, as assessed by a DNMT task, but possibly interferes with more subtle aspects of memory-guided decision processes.

## Acknowledgements

We are indebted to IMM’s Bioimaging, Comparative Pathology, and Rodent facilities for critical help. We want to thank all the members of the Remondes Lab for fruitful discussions and help.

## Funding reference

UID/BIM/50005/2022, project funded by Fundação para a Ciência e a Tecnologia (FCT)/ Ministério da Ciência, Tecnologia e Ensino Superior (MCTES) through Fundos do Orçamento de Estado.

FCT granted a PhD Fellowship (PD/BD/128107/2016, COVID/BD/151601/2021) to ASC, and an Exploratory Grant (IF/00201/2013), an IMM Director’s Fund Award, an Investigator FCT Position at IMM-JLA (IF/00201/2013), a Research Grant (PTDC/MED-NEU/29325/2017), and a CEEC 2022 2022.00811.CEECIND Principal Investigator contract to MR.

## METHODS

### LEAD CONTACT AND MATERIALS AVAILABILITY

Further information and requests for resources and reagents should be directed to and will be fulfilled by Lead Contact, Miguel Remondes, DVM, PhD. This study did not generate new unique reagents.

### EXPERIMENTAL MODEL AND SUBJECT DETAILS

We used 17 male Long-Evans rats, purchased from Charles River Europe, aged between 12 to 24 weeks old at the start of the experiment kept in a 12hr light/dark cycle, fed *ad libitum*, housed in pairs until surgeries, and single-caged thereafter. All procedures were performed in accordance with EU and Institutional guidelines.

### BEHAVIORAL APPARATUS

The behavioral apparatus consisted of (1) a T-maze, with: a 25 cm x 29.5 cm start region; a 170 cm long central arm; a 170 cm long top arm and two 64.5 cm long side arms. All arms had a 10 cm width and 5 cm walls, (2) a resting box (ITI) 57×39×42 cm. Two reward ports (2 cm diameter), connected by tubes to two peristaltic pumps, were placed at the end of the side arms. A cable tray, placed over the maze, supported: (1) a Flea 3 PointGrey^TM^ camera, used to record the behavioral sessions at a rate of 30 fps; (2) the optogenetic light delivery system. Both were connected to a computer, in an adjacent chamber outside the behavior room. Video recording, position tracking, reward delivery and optogenetic stimulation onset were processed and/or controlled by Bonsai software (Lopes et al., 2015).

Optogenetic light delivery was achieved using the PlexBright system (Plexon, Dallas, TX, USA). One LD-1 Single Channel LED Driver, set at 240 mA, was connected to a Dual LED Commutator, placed above the maze. The commutator was connected to two Orange 620 nm LED modules, with each attached to a 2 meters-long 0.66 NA patch cable with a FC ferrule tip and stainless-steel cladding. The patch tip was connected to an implanted lambda fiber stub (200/230 um, 0.66 NA), with a 2mm active length and 5 mm implant length (Optogenix^TM^), through a ceramic sleeve. Depending on the number of fiber stubs implanted (please see below), either one or two patch cables were used. Coupled to the tip of the patch cable and close to the implant, a red 5 mm LED was used to track the animal’s xyt position coordinates.

### VIRAL INJECTIONS AND STUB IMPLANTS

The AAV9-hSyn-CTRLeYFP and the AAV9-hSyn-eYFP were purchased from Addgene^TM^ Plasmid Repository (Watertown, MA, USA). Viral titers were 1×10^13^ vg/mL and 7×10^12^ vg/mL, respectively. Vector suspensions were micro-injected bilaterally in the CG region, as described below.

Rats were anesthetized and prepared for craniotomy as previously published (Ferreira-Fernandes et al., 2019). The skin was then incised and retracted to expose the skull, four anchoring screws were firmly secured on the laterals of the parietal bone around the injection sites, where a craniotomy and durotomy were performed. The tip of a glass micro-pipette loaded with the tracer was lowered into the brain at the following coordinates: A/P: +1 mm, M/L: +/− 0.5 mm, D/V: −1.6 mm.

Following the injections, a fiber stub was mounted on the frame, using an in-house custom connector. In 6 rats (5 eNpHR3.0+ and 1 CTRL) two fiber stubs were implanted at A/P: +1 mm, M/L: +/− 1 mm, D/V: - 1 mm, at a 15-degree angle. In the remaining 11 rats, only one diagonally oriented (30-degree angle) fiber stub was implanted, on the rat’s right hemisphere, such that it crossed the CG region bilaterally, at A/P: +1 mm, M/L: + 1 mm, D/V: −2.8 mm. The fiber stubs were secured in place with dental acrylic (DA) connecting them to the anchoring screws. Once DA was cured, the skin was sutured, and a 10% povidone iodine solution was applied in the skin around the wound. Carprofen solution (½ dose) was again injected subcutaneously. The wound was sutured, and the rat was allowed to recover. Overall, 7 CTRL and 10 eNpHR3.0+ animals were used.

### BEHAVIORAL TASK AND PROTOCOL

For one week before surgery, rats were handled by the experimenter for ten minutes daily and offered chocolate milk, through a 1 mL syringe. This was repeated one week after post-surgical recovery. On the first day in the behavioral apparatus rats were allowed to freely explore it for 10 minutes, with rewards available in both ports. On subsequent sessions rats performed the DNMT protocol for a maximum of 30 trials per session and one session per day.

The delayed non-match to position (DNMP) protocol is a hippocampus-dependent task (Aggleton et al., 1986; Racine and Kimble, 1965; Wood et al., 2000) typically used to study working memory in rodents (Dudchenko, 2004). The task is run on a T-shaped maze, where a central stem arm leads to two choice arms (left and right) at whose end reward is delivered as appropriate. Each trial was composed of three epochs: (1) the sample run; (2) the delay; (3) the test run. In the sample run, access to one of the choice arms was blocked and the animal is forced to turn to the only available arm where it received a reward. This was followed by a 30 sec delay period, with the rat placed outside the maze, after which the rat performs the choice run. In the choice run the rat is placed back on the central arm, allowed free access to both choice arms, and rewarded for choosing the previously blocked arm, e.g. for choosing a non-match to position arm. An ITI period followed, with the rat outside the maze while the experimenter prepared the following trial, never surpassing 30 seconds. Each arm was randomly assigned the role of “Sample” for a maximum of three consecutive trials. When rats reached a performance level of at least 75% the patch cable was connected to the fiber stub(s) at the beginning of each behavioral session.

### ILLUMINATION PROTOCOL

The illumination protocol started once rats reached 75% performance for three consecutive sessions. It consisted in three illumination conditions depending on the trial epoch being illuminated: (1) No illumination, (NI); (2) Sample illumination (SI), in which the sample run was illuminated; (3) Test illumination (TI), in which the test run was illuminated. Illumination lasted for 15 seconds. Light onset was triggered by the rat’s entrance into the central arm. The three illumination conditions were interleaved, balanced, and pseudo-randomized across trials and within session. We allowed no more than three consecutive trials with the same illumination condition. The full illumination protocol lasted for 15 sessions, corresponding to the collection of 150 trials per illumination condition. In two eNpHR3.0+ rats, we only collected 40 and 50 per condition, respectively, in one CTRL rat we collected no SI trials.

At the end of the experiment, rats were sacrificed, and the implant was retrieved and connected to the light delivery system for visual confirmation of light output through the fiber stub. Rats were then sacrificed with an overdose of sodium pentobarbital (800 mg/kg) and transcardially perfused with 250 mL of PBS, followed by 250 mL of 10% neutral buffered formalin. Their brains were removed and stored in a Falcon tube containing 40 mL of 10% neutral buffered formalin, covered with aluminum foil. The brain samples were changed to a 15% sucrose solution, and kept at 4°C overnight. They were then embedded in gelatin, frozen in 2-Methylbutane liquid nitrogen and cut into 50 µm coronal sections, at −25°C, using a cryostat (Leica, CM3050 S), from A/P +4 mm to −2 mm. For DAPI staining, the gelatin was removed, the slices were washed and incubated with DAPI (1:700) at room temperature for 20 minutes. After washing they were mounted in slides with Mowiol^TM^, and left to dry for 24 hours.

Brain slices were imaged using a Zeiss AxioZoom V16 fluorescence stereo microscope, equipped with a AxioCam MRm (CCD, 1388×1040 px) camera and a 1x PlanNeoFluar Z (.25 NA) objective. A Zeiss GFP filter set (FS83HE) was used to observe CTRLeYFP or eYFP and a Zeiss DAPI filter set (FS01) was used to observe DAPI stained nuclei.

### QUANTIFICATION AND STATISTICAL ANALYSIS

Each trial was classified as a correct or incorrect trial. Each session’s performance was calculated as the number of correct trials, over the total number of trials in the session. Performance under each illumination condition was calculated as the number of correct trials over the total number of trials for that condition. Rat’s xy position was recorded and timestamped, subdivided into individual runs, and labelled according to run type (sample or test run), illumination condition and outcome (correct or incorrect trial). We then calculated the latency to choice point and the time spent in choice point (Fig. S2) for each test run. The latency to choice point was calculated as the difference between the timestamp collected upon entering the choice point and the timestamp collected upon entering the central arm. The time spent in choice point was calculated as the difference between the timestamps of choice point exit and entry. The choice point was determined as the measured limits of the choice point square on the maze, to which an extra 10 cm were added (Fig. S2).

Since light delivery had a maximum duration of 15 seconds, runs in which the sum of the latency and time spent in choice point was higher than 15 seconds were removed from the analysis (563 trials, around 7% of the total number of trials).

Custom made python scripts were used for all data handling and metric computations. In all the analyses conducted we resorted to generalized linear mixed models with individual rat as a random factor. The selection of random terms was achieved by considering the experimental design, namely by adding random effect terms that could vary within rat / subject, and the lowest Akaike Information Criterion (AIC). To determine the effect of illumination condition, group, session number, trial number and their interactions on trial outcome we used a Binomial Distribution and a logit link function. To determine the effect of the previous variables and outcome on latency to the choice point and time at the choice point we tested the Gamma and Inverse Gaussian Distributions with an identity or inverse link functions, in a total of four combinations. The distributions and link functions to test reaction time variables were selected as recommended in Lo and Andrews, 2015 (Lo and Andrews, 2015). The choice of both was determined by model convergence (or lack of) and the lowest AIC. This resulted in a Gamma distribution with an inverse link function for latency to choice point, and the Inverse Gaussian distribution with an identity link function for time spent at choice point. An omnibus (Wald) Chi-Squared test was used to test the main effects of the independent variables and their interaction. *Post-hoc* multiple comparisons were performed on the estimated marginal means extracted from the models. All this analysis was conducted using Jamovi with the GAMLJ package. *Pos-hoc* comparisons were also conducted in Jamovi 2.3, with the emmeans package (Jonathon Love, 2022).

