## Supplementary material for "Effects of optogenetic silencing the Anterior Cingulate Cortex in a delayed non-match to trajectory task": Figure S1 (from another version of this preprint: version 2, 2023-09-06)

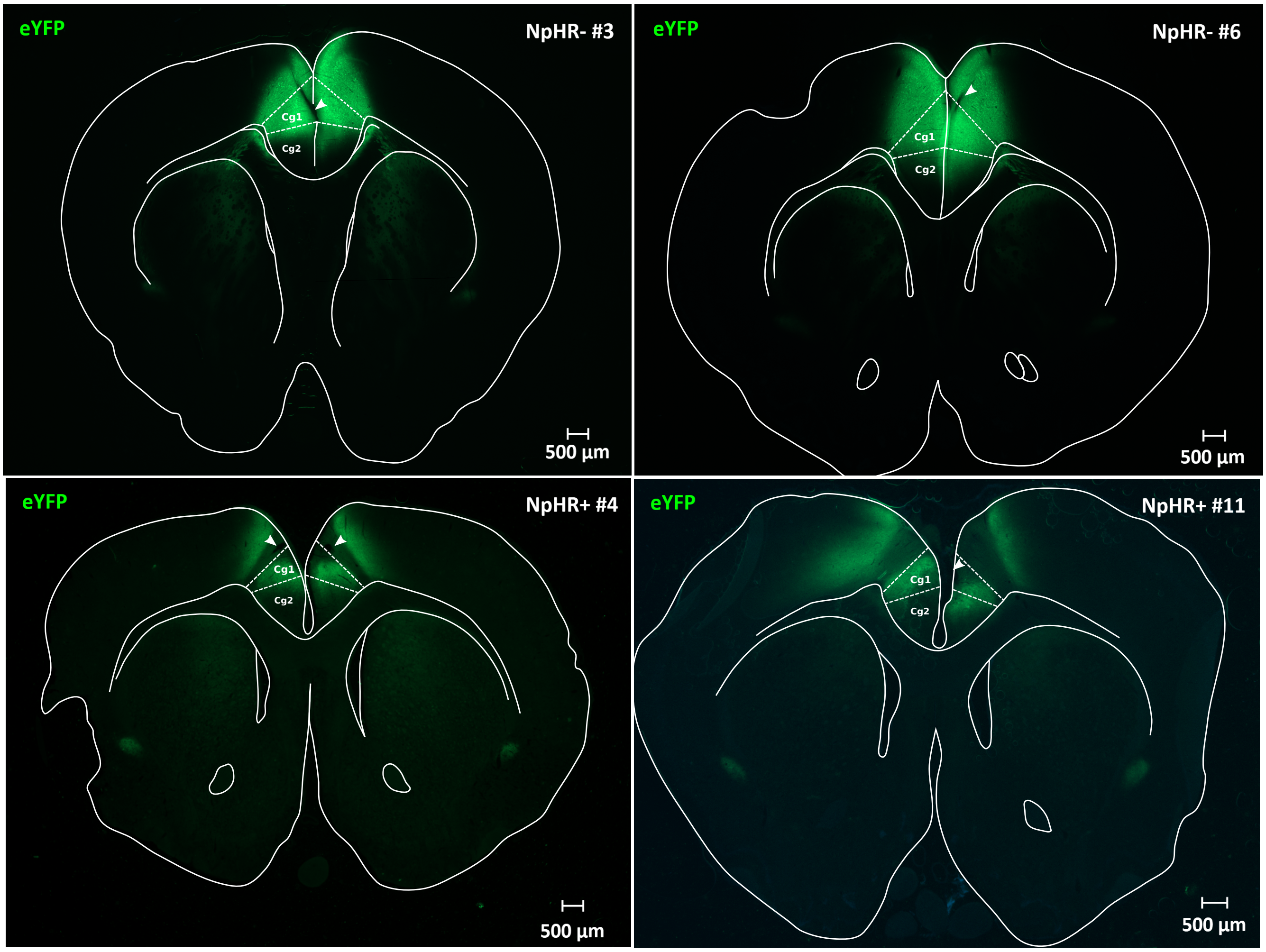


**Figure S1. Histological verification of CTRLeYFP or eYFP expression after bilateral injection of viral constructs.** (A) Examples eYFP expression and fiber entry lesion (arrow) in CTRL rats. (B) Similar to previous but for eYFP in eNpHR3.0+ rats: with two (left) or only one (right) fibers implanted. CG regions were outlined as in Paxinos Watson Rat Brain Atlas, at +1 A/P.
