## Supplementary material for "Effects of optogenetic silencing the Anterior Cingulate Cortex in a delayed non-match to trajectory task": Figure S2 (from another version of this preprint: version 2, 2023-09-06)

**A**

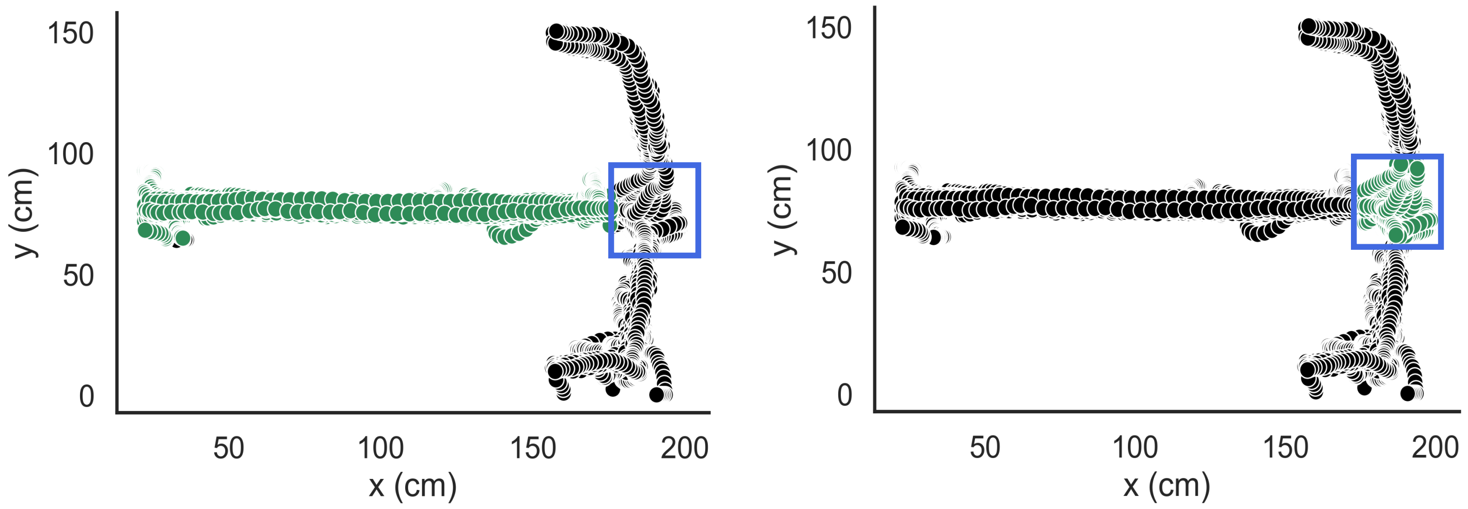


**B**

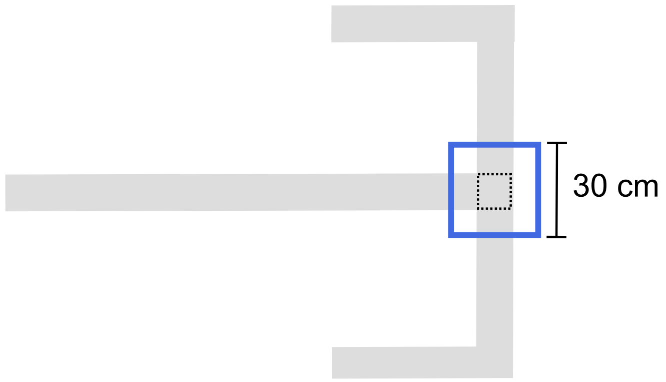


**Figure S2. Measurement of latency to choice point and time spent in choice point.** (A) Example position data collected from one session. Left, the position data before entering the choice point is colored green. Right, the position data inside the choice point is colored green. Latencies were computed from the timestamps corresponding to each data point series. (B). Scheme of the choice area used for latency calculations (in blue) corresponding to and the actual maze choice point (black square) plus 10 cm around.
